# Genetic depletion studies define receptor usage by virulent hantaviruses in human endothelial cells

**DOI:** 10.1101/2020.12.30.424861

**Authors:** M. Eugenia Dieterle, Carles Solà-Riera, Rohit K. Jangra, Chunyan Ye, Samuel M. Goodfellow, Eva Mittler, Ezgi Kasikci, Steven B. Bradfute, Jonas Klingström, Kartik Chandran

**Affiliations:** Department of Microbiology and Immunology, Albert Einstein College of Medicine, Bronx, NY 10461, USA; Center for Infectious Medicine, Department of Medicine Huddinge, Karolinska Institutet, 141 86 Stockholm, Sweden; University of New Mexico Health Science Center, Center for Global Health, Department of Internal Medicine, Albuquerque, NM 87131, USA

## Abstract

Hantaviruses are a large group of RNA viruses that include known epidemic threats and other agents poised for emergence. Several rodent-borne hantaviruses cause zoonoses accompanied by severe illness and death. However, assessments of zoonotic risk and the development of countermeasures alike are challenged by our limited knowledge of the molecular mechanisms of hantavirus infection, including the identities of cell entry receptors and their roles in influencing viral host range and virulence. Previous work has implicated several cell-surface molecules, most notably β3- and β1-containing integrin heterodimers, decay-accelerating factor (DAF), and the cadherin superfamily protein protocadherin-1 (PCDH1), in hantavirus entry in endothelial cells, the major targets of viral infection in humans. Despite the fact that β3/β1 integrins have been presumed to be the major hantavirus entry receptors for over two decades, rigorous genetic evidence supporting their requirement, and that of DAF as an entry cofactor, is lacking. Here, we used CRISPR/Cas9 engineering to knock out four candidate hantaviral receptors, singly and in combination, in a human endothelial cell line that recapitulates the properties of primary microvascular endothelial cells. PCDH1 loss substantially reduced entry and infection by a subset of hantaviruses endemic to the Americas. In contrast, the loss of β3 integrin, β1 integrin, and/or DAF had little or no effect on entry by any of a large panel of hantaviruses tested. We conclude that the major host molecules necessary for endothelial cell entry by PCDH1-independent hantaviruses remain to be discovered.

---

Hantaviruses are a large group of enveloped viruses with segmented negative-strand RNA genomes whose members infect a wide range of mammalian hosts (1). The zoonotic transmission of some rodent-borne hantaviruses is associated with two major diseases in humans, hemorrhagic fever with renal syndrome (HFRS) and hantavirus cardiopulmonary syndrome (HCPS), in endemic regions of Europe/Asia and North/South America, respectively. Most HFRS cases are associated with the ‘Old-World hantaviruses’ Hantaan virus (HTNV), Seoul virus (SEOV), Dobrava-Belgrade virus (DOBV) and Puumala virus (PUUV), whereas HCPS is primarily caused by the ‘New-World hantaviruses’ Andes virus (ANDV) and Sin Nombre virus (SNV). Although humans are typically dead-end hosts for rodent-borne hantaviruses, several incidents of ANDV person-to-person transmission, including a recent superspreader event, have been documented, underscoring the epidemic threat posed by hantaviruses (2, 3).

Integrins (αVβ3, α5β1, αMβ2 and αXβ2) and components of the complement system (e.g., decay accelerating factor, DAF) have been proposed as candidate hantavirus entry factors (4–8). β3- and β1-containing integrins have been presumed to be the major receptors for virulent and avirulent hantaviruses, respectively, for over two decades (5, 9, 10). More recently, we identified protocadherin-1 (PCDH1) as a critical entry determinant for New-World hantaviruses (11). However, evidentiary support for each of the above putative receptors differs considerably. *PCDH1* was identified in a comprehensive genetic screen for ANDV entry factors. Its genetic depletion in human microvascular endothelial cells, the major targets of infection *in vivo*, inhibited viral entry and infection. Further, genetic ablation of *PCDH1* in Syrian hamsters reduced ANDV multiplication and protected them from lethal HCPS-like disease (11). By contrast, none of the other candidate receptors were observed as hits in two independent genetic screens for hantavirus host factors (11–13). To our knowledge, their requirement has also not been rigorously evaluated via genetic approaches in physiologically relevant *in vitro* and *in vivo* models.

Here, we used a genetic depletion/complementation strategy to investigate hantavirus receptor requirements in a human microvascular endothelial cell line, TIME, a genetically manipulable model for primary endothelial cells (14). We first confirmed that TIME cells resembled primary human umbilical vein endothelial cells (HUVEC)–the standard model for *in vitro* hantavirus studies–in expressing key endothelial surface markers as well as the candidate hantavirus receptors (Fig 1a). Further, TIME cells were susceptible to hantavirus entry mediated by divergent Gn/Gc proteins (Fig 2), and ANDV Gn/Gc-dependent entry in these cells was selectively inhibited by a PCDH1-specific monoclonal antibody, mAb-3305 (Fig 2a).

**Figure 1.**
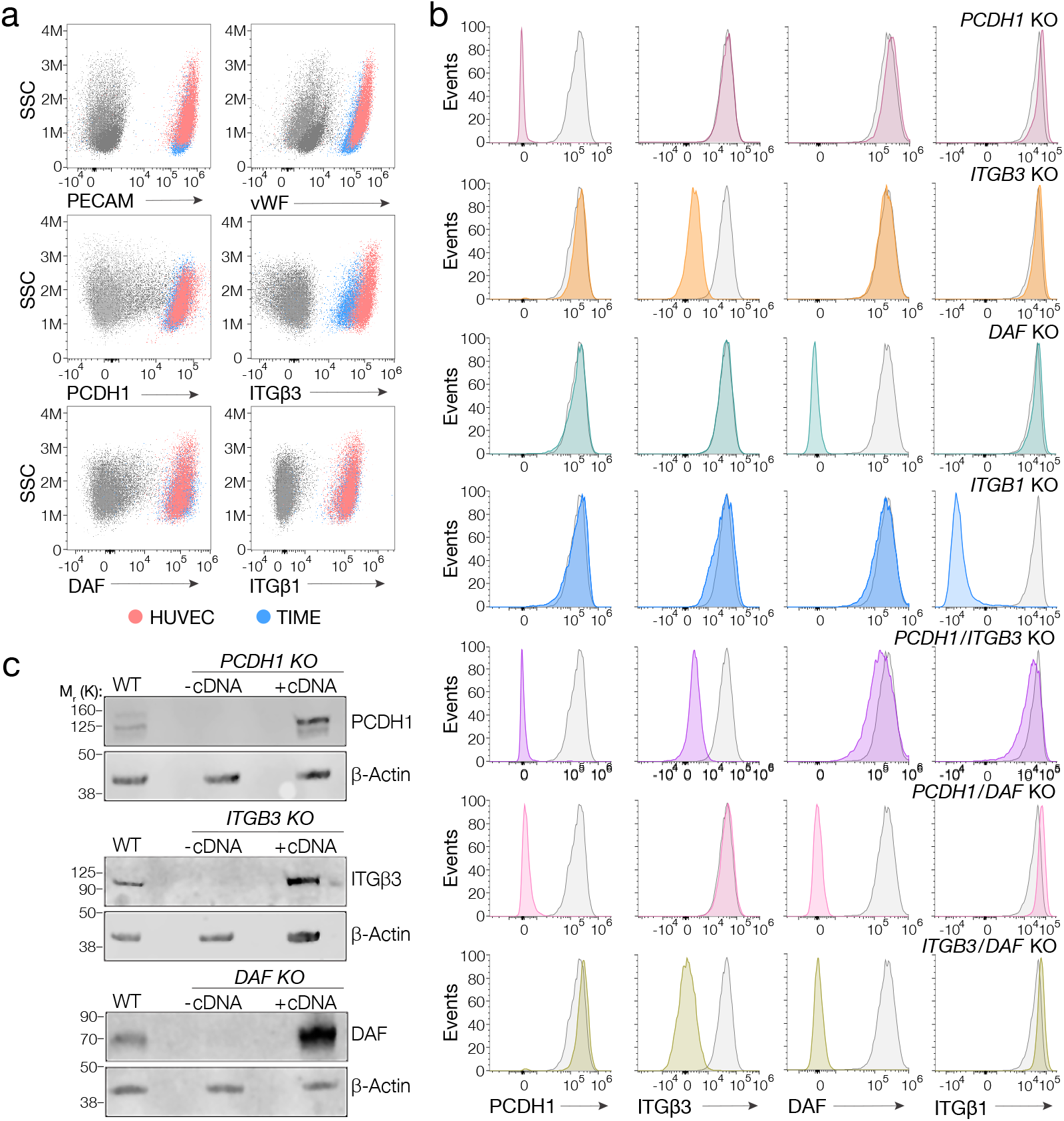
Suitability of TIME cells as a model to study hantavirus entry and the generation of knockout cells. **(a)** Upper panels, total flow cytometry plots of HUVEC and TIME cells stained for endothelial cell markers PECAM and vWF. Medium and lower panels, surface flow cytometry plots of HUVEC and TIME cells stained for the indicated markers. **(b)** Surface flow cytometry of WT and KO TIME cells stained as above. Histograms of WT cells are shown in gray; single- and double-KO cells are shown in color. **(d)** Western blot analysis of WT TIME cells and KO cells ± cDNA. β-Actin was used as a loading control.

**Figure 2.**
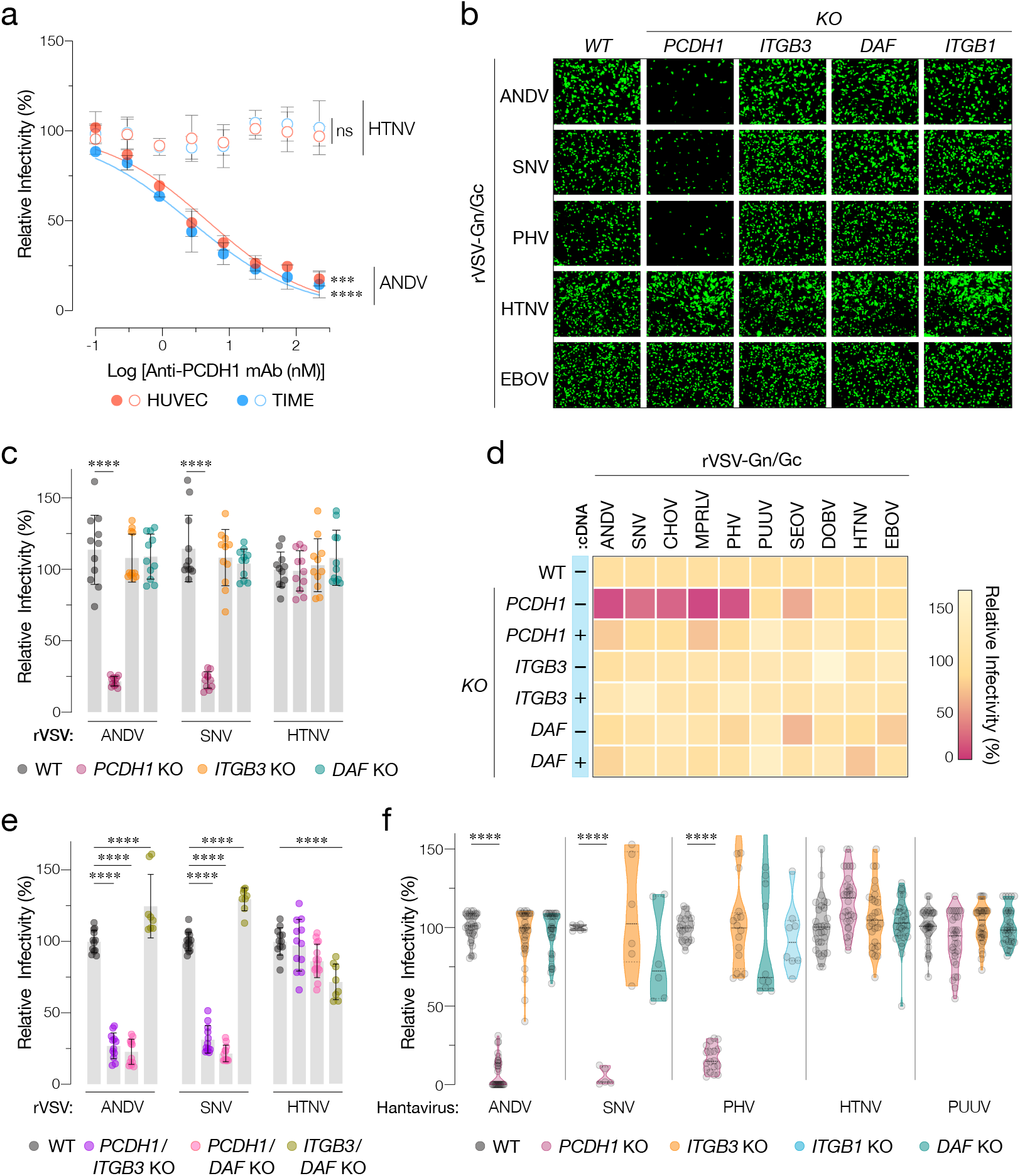
Hantavirus receptor requirement in endothelial cells. **(a)** Capacity of PCDH1 monoclonal antibody (mAb-3305) to block viral entry. HUVEC and TIME cells were preincubated with mAb-3305, and then exposed to the indicated rVSVs. TIME cells, n=6 from 3 independent experiments, HUVEC, n=4 from 2 independent experiments. **(b)** Representative images of eGFP-positive rVSV-infected WT and KO TIME cells. **(c)** WT and KO cells were exposed to the indicated rVSV-Gn/Gc. n=11 for each cell line from 3 independent experiments. **(d)** WT and KO cells ± corresponding cDNA were exposed to rVSV-Gn/Gc. Viral infectivities (averages from 3 independent experiments) are shown in the heatmap. **(e)** WT and double-KO cells were exposed to rVSV-Gn/Gc. *PCDH1/ITGB3* KO, n=12; *PCDH1/DAF* KO, n=12; and *ITGB3/DAF* KO, n=9 from 3 independent experiments. **(f)** Cells were exposed to authentic hantaviruses and infected cells were manually enumerated by immunofluorescence microscopy for ANDV, HTNV, and PUUV (each point represents infectivity of the average of positive cells per field relative to WT). Data are from 2 independent experiments. PHV and SNV-infected cells were detected and enumerated by automated imaging following immunofluorescence staining. For PHV, WT and *PCDH1* KO n=18, *ITGB3* KO n=16 from 4 independent experiments; *DAF* KO, n=10, *ITGB1* KO n=8, from 3 independent experiments. For SNV WT, *ITGB3* KO, *DAF* KO n=6; *PCDH1* KO n=5 from 3 independent experiments. Averages±SD are shown. WT vs. KO comparisons by two-way ANOVA with Tukey’s test (a, f) or Dunnett’s test (c, e) ***, P<0.0002; ****, P < 0.0001. Other comparisons were not significant (P>0.05, ns).

We used CRISPR/Cas9 genome engineering to knock out (KO) *PCDH1*, *DAF*, and the genes encoding β3 and β1 integrins (*ITGB3* and *ITGB1*) in TIME cells. Loss of specific protein expression at the cell surface and in cell extracts was verified by flow cytometry and western blotting, respectively (Fig 1b–c). We next exposed WT and KO TIME cells to recombinant vesicular stomatitis viruses carrying the Gn/Gc glycoproteins of ANDV, SNV, and HTNV (rVSV-Gn/Gc) (Fig 2b–d). An rVSV bearing the Ebola (EBOV) virus glycoprotein was used as a negative control. *PCDH1* KO substantially diminished ANDV and SNV Gn/Gc-mediated infection but had no discernible effect on infection by HTNV Gn/Gc. Unexpectedly, *ITGB3* and *DAF* KOs did not inhibit cell entry mediated by any of these glycoproteins (Fig 2b–c). We extended these studies to rVSVs bearing Gn/Gc proteins from the New-World hantaviruses Choclo virus (CHOV), Maporal virus (MPRLV), Prospect Hill virus (PHV), and the Old-World hantaviruses PUUV, SEOV, and DOBV (Fig 2d). Loss of *PCDH1* substantially reduced cell entry mediated by the glycoproteins from the New-World viruses but not the Old-World viruses, in a manner that could be restored by complementation with *PCDH1* cDNA. By contrast, none of these viruses displayed a requirement for *ITGB3* or *DAF*. These findings confirm and extend the critical role played by PCDH1 in cell entry by New-World hantaviruses. However, they also contradict the claims that β3 integrin and DAF are necessary for hantavirus entry in endothelial cells.

To account for potential receptor redundancy and cross-talk, we next evaluated double-KO populations of TIME cells (Fig 1b). *PCDH1/ITGB3* and *PCDH1/DAF* KO cells resembled *PCDH1* KO cells in their susceptibility to rVSV-Gn/Gc infection (Fig 2e), indicating that the loss of PCDH1 did not unmask viral requirements for β3 integrin or DAF, or vice versa. Further, *ITGB3/DAF* KO cells remained fully susceptible to viral entry (ANDV, SNV) or subject to only a modest (~25%) reduction (HTNV) (Fig 2e). Thus, the lack of viral entry phenotypes in *ITGB3* and *DAF* single-KO cells cannot be explained by the functional redundancy of these two proteins.

Finally, we evaluated the single-KO TIME cells for infection with authentic hantaviruses by immunofluorescence microscopy (Fig 2f), which corroborated our findings with rVSVs. Specifically, we observed a critical role for PCDH1 in endothelial cell infection by the New-World hantaviruses ANDV, SNV, and PHV, and no apparent roles for β3 integrin and DAF in entry by virulent New-World and Old-World hantaviruses. Concordantly, quantitative real-time PCR revealed a substantial reduction in the generation of SNV progeny genomes at 24 h post-infection in cells deficient for PCDH1 (5±2%) but not β3 integrin (108±15%) or DAF (82±13%; mean±SEM, n=6 from two independent experiments for each cell line). Moreover, and contrary to an earlier report (5), we also found no apparent role for β1 integrin in entry by the avirulent New-World hantavirus PHV.

Our results underscore PCDH1’s critical role as an entry receptor for New-World hantaviruses in endothelial cells but indicate that three other major candidate receptors (β3 integrin, β1 integrin, DAF) are dispensable. We note that our results do not rule out that one or more of these proteins are involved in hantavirus entry in other cell types or that they are involved in endothelial cell subversion post-viral entry. The sum of the evidence does indicate, however, that the major host molecules necessary for entry and infection in endothelial cells by the PCDH1-independent, HFRS-causing Old-World hantaviruses remain to be discovered.

## Materials and methods

### Generation of KO cell populations

Gene-specific single-guide RNAs were cloned into lentiCRISPR v2 and CRISPR-Cas9 engineering was performed as described (11). Receptor-negative subpopulations of TIME cells were isolated by FACS-sorting. The targeted genomic loci in genomic DNA isolated from these subpopulations were verified by Sanger sequencing. All sequenced clones bore a frameshift inducing-indel at the targeted site.

### rVSVs and infections

rVSVs expressing a fluorescent reporter and hantavirus Gn/Gc were described previously (11, 12, 15) or generated as described (12). Viral infectivity was measured by automated enumeration of fluorescent cells (12). 100% relative infectivity corresponds to 18–35% infected cells for ANDV, SNV, HTNV, EBOV, CHOV, MPRLV; 8–15% for SEOV, DOBV, PHV and PUUV.

### Hantaviruses and infections

ANDV isolate Chile-9717869, HTNV isolate 76-118, PUUV isolate Sotkamo, PHV, and SNV isolate SN77734 were used. Cells were infected with the respective virus. After 48 h (24 h for SNV), cells were immunostained (16). Cells were imaged and enumerated by fluorescence microscopy (11, 16).

Extended methods are available for this paper.

## Acknowledgments

This work was supported by NIH grant R01AI132633 (to K.C.), and Swedish Research Council grant 2018-02646 (to J.K). M.E.D. was partially supported as a Latin American Fellow in the Biomedical Sciences of the Pew Charitable Trusts. S.M.G. was partially supported by the UNM HSC Infectious Disease and Inflammation Program NIH grant T32AI007538. The Einstein Flow Cytometry Core is partially supported by NIH grant P30CA013330.

## Extended Methods

### Cells

HUVEC (C2517A-Lonza) and TIME cells (ATCC CRL4025) were cultured as described (1, 2).

### Surface and total flow cytometry

Antibodies: AF480-Rabbit-α–vWF (ab195028-Abcam), PeCy7-Mouse-α–PECAM (563651-BD), APC-Mouse-α–β1-Integrin (559883-BD), AF 647-Mouse-α–β3-Integrin (336407-Biolegends), PE-Mouse-α–DAF (555694-BD); α–PCDH1 mAb 3305 (2) /AF488-α–Human (A-11013-Invitrogen). For surface staining, cells were kept at 4°C. Foxp3/Transcription Factor Staining Buffer Kit was used for intracellular staining following the manufacturer’s instructions (TNB-0607-KIT).

### Western blotting

Performed as described (2). Antibodies: Mouse α–PCDH1 (sc-81816-Santa Cruz) 1:200; Rabbit-α–β3 Integrin (#4702-Cell Signaling) (1:300); Mouse-α–DAF (NaM16-4D3-Santa Cruz) 1:200; Mouse α–β-Actin (sc-4778-Santa Cruz) (1:300). IRDye 680LT Goat α-Rabbit IgG 680 or IRDye 680LT Goat α-Mouse secondaries Abs (LI-COR, Lincoln, NE) were used at a dilution of 1:10,000, and the final blot was then imaged using a LI-COR Fc fluorescent imager.

**mAb-3305 mediated inhibition of virus entry** as described earlier (2). Briefly, HUVEC and TIME cells were preincubated with mAb 3305 (0-222 nM), and then exposed to rVSVs bearing ANDV or HTNV Gn/Gc. NH_4_Cl (20 mM) was added 1 hour after infection. Viral infectivity was measured 14 hours post infection by automated enumeration of eGFP positive cells using a Cytation5 automated fluorescence microscope (BioTek) and analyzed using the Gen5 data analysis software (BioTek).

### Generation of KO cell populations

A CRISPR sgRNA was designed to target, *PCDH1* (2); *ITGB3* 5′-<u>CCA</u>CGCGAGGTGTGAGCTCCTGC-3’; *DAF* 5′-<u>CCC</u>CCAGATGTACCTAATGCCCA-3 and *ITGB1* 5′-ATACAAGCAGGGCCAAATT<u>GTGG</u>-3’ (protospacer acceptor motif [PAM] is underlined). sgRNAs were cloned into lentiCRISPR v2 (Addgene plasmid # 52961) and CRISPR-Cas9 engineering was performed as described (2). Receptor-negative subpopulations of TIME cells were isolated by FACS-sorting. The targeted genomic loci in genomic DNA isolated from these subpopulations were amplified by PCR, and the amplicons were TA-cloned into the pGEM-T vector. 15–20 clones for each KO cell population were subjected to Sanger sequencing. For each KO cell population, all sequenced clones showed an indel at the targeted site, resulting in a frameshift that brought one or more stop codons into frame.

### rVSVs and infections

rVSVs expressing eGFP and bearing Gn/Gc from ANDV, SNV, HTNV, SEOV, DOBV, MPRLV, EBOV, PHV, PUUV were described previously (2–6). rVSV expressing mNeon Green fused to the VSV phosphoprotein and bearing Gn/Gc from CHOV (GenBank accession number KT983772.1) was engineered, rescued and propagated as described before (4). Viral infectivity was measured 14 hours post infection by automated enumeration of eGFP- or mNeongreen- positive cells using a Cytation5 automated fluorescence microscope (BioTek) and analyzed using the Gen5 data analysis software (BioTek). 100% relative infectivity corresponds to 18-35% infected cells for ANDV, SNV, HTNV, EBOV, CHOV, MPRLV. 8-15% for SEOV, DOBV, PHV, PUUV.

### Hantavirus and infections

ANDV isolate Chile-9717869, HTNV isolate 76-118, PUUV isolate Sotkamo, PHV, and SNV isolate SN77734 were used. TIME cells were seeded in 24-well plates (on glass coverslips) or 96 well plate, and infected with a multiplicity of infection of 1. After 48 hours (24 hs for SNV), immunofluorescence was done as described before (7). Briefly, human polyclonal antibodies from convalescent patient serum plus anti-hantavirus nucleocapsid protein B5D9 monoclonal antibody (Progen) were used for HTNV, PUUV, ANDV. Rabbit polyclonal sera specific for SNV nucleoprotein NR-12152 (BEI Resources) was used for PHV and SNV and incubated for an hour. For HTNV, PUUV, ANDV infected cells, AF488-α–Human and α-Mouse IgG antibodies (Life Technologies) were used as secondaries. For SNV and PHV, AF488-α–Rabbit IgG was used (Life Technologies). Counterstaining of nuclei was done with DAPI (Thermo Fisher Scientific). Cells were imaged by fluorescence microscopy (Nikon, Eclipse TE300) with a 60X objective or by automated enumeration of eGFP-positive cells using a Cytation5 automated fluorescence microscope (BioTek) as described.

### SNV RNA levels

WT and KO TIME cells were harvested before infection (negative control) or at 24 hours after infection. Degenerate primers and probe were adopted from (8) based on the S-segment of the SNV genome. Human β-Actin primers with a VIC™/TAMRA™-dye probe (Applied Biosystems, Cat. # 4310881E) was used as an endogenous control. The qPCR reactions were carried out by using TaqMan™ Fast Advanced Master Mix (Applied Biosystems, Thermo Fisher Scientific) following instruction manual. Relative gene fold change was calculated by normalizing SNV to β-actin in cells using ΔΔCt values. Averages ±SEM from two independent experiments are shown.

### Statistics and reproducibility

The number of independent experiments and the measures of central tendency and dispersion used in each case are indicated in the figure legends. The testing level (alpha) was 0.05 for all statistical tests. Statistical comparisons were carried out by two-way ANOVA with a post hoc correction for family-wise error rate. All analyses were carried out in GraphPad Prism.

